# Exome-wide association studies in general and long-lived populations identify genetic variants related to human age

**DOI:** 10.1101/2020.07.19.188789

**Authors:** Patrick Sin-Chan, Nehal Gosalia, Chuan Gao, Cristopher V. Van Hout, Bin Ye, Anthony Marcketta, Alexander H. Li, Colm O’Dushlaine, Dadong Li, John D. Overton, Jeffrey D. Reid, Aris Baras, Regeneron Genetics Center, David J. Carey, David H. Ledbetter, Daniel Rader, Marylyn D. Ritchie, Scott M. Damrauer, Sofiya Milman, Nir Barzilai, David J. Glass, Aris N. Economides, Alan R. Shuldiner

**Author notes:** Corresponding author: Alan R. Shuldiner, MD, 777 Old Saw Mill River Road, Tarrytown, New York 20591.

## Abstract

Aging is characterized by degeneration in cellular and organismal functions leading to increased disease susceptibility and death. Although our understanding of aging biology in model systems has increased dramatically, large-scale sequencing studies to understand human aging are now just beginning. We applied exome sequencing and association analyses (ExWAS) to identify age-related variants on 58,470 participants of the DiscovEHR cohort. Linear Mixed Model regression analyses of age at last encounter revealed variants in genes known to be linked with clonal hematopoiesis of indeterminate potential, which are associated with myelodysplastic syndromes, as top signals in our analysis, suggestive of age-related somatic mutation accumulation in hematopoietic cells despite patients lacking clinical diagnoses. In addition to *APOE*, we identified rare *DISP2* rs183775254 (p = 7.40×10^−10^) and *ZYG11A* rs74227999 (p = 2.50×10^−08^) variants that were negatively associated with age in either both sexes combined and females, respectively, which were replicated with directional consistency in two independent cohorts. Epigenetic mapping showed these variants are located within cell-type-specific enhancers, suggestive of important transcriptional regulatory functions. To discover variants associated with extreme age, we performed exome-sequencing on persons of Ashkenazi Jewish descent ascertained for extensive lifespans. Case-Control analyses in 525 Ashkenazi Jews cases (Males ≥ 92 years, Females ≥ 95years) were compared to 482 controls. Our results showed variants in *APOE* (rs429358, rs6857), and *TMTC2* (rs7976168) passed Bonferroni-adjusted p-value, as well as several nominally-associated population-specific variants. Collectively, our Age-ExWAS, the largest performed to date, confirmed and identified previously unreported candidate variants associated with human age.

## INTRODUCTION

Over the past decades, the average human life expectancy has increased dramatically, which has resulted in more people living into adulthood and older ages (WHO, 2015). The number of people in the United States aged 65 years or older is expected to double from ∼43.1 million in 2012 to ∼83.7 million in 2050 (Ortman et al., 2014). However, increases in life expectancy have not been accompanied by equivalent increases in disease-free lifespan or ‘healthspan’. Both the Center for Disease Control and World Health Organization report that the leading causes of death in seniors are cardiovascular diseases, malignancies and neurodegenerative disorders, which suggests aging as the strongest risk factor from developing these age-associated pathologies (Heron, 2019; WHO, 2015). A deeper understanding of aging biology could lead to interventions to decrease the incidence of these age-related diseases, thus increasing healthspan.

There are many factors that may influence human lifespan, which includes diet, gender, ancestry, public health, education, family life, socioeconomic status, social responsibility, access to medical care and genetics. Indeed, several studies have shown a strong familial component to living to older ages. For example, siblings of centenarians exhibit greater chances of living to the ages of their oldest sibling, as compared to the general population (Sebastiani et al., 2016). Moreover, the offspring of long-lived individuals (LLI) are more likely to exhibit delayed onset of age-related diseases and compressed disease morbidity (Atzmon et al., 2004; Dutta et al., 2013; Dutta et al., 2014; Gudmundsson et al., 2000; Lipton et al., 2010; Terry et al., 2003). Multiple reports indicate a genetic component to aging, in which current literature suggests heritability to be between ∼20-35% (Finch and Tanzi, 1997; Herskind et al., 1996; Sanders et al., 2014; van den Berg et al., 2017), although more recent studies on millions of participants predict lifespan heritability to be much lower (Kaplanis et al., 2018; Timmers et al., 2019). Notably, heritability of lifespan increases at older ages, where effects are most prominent in centenarians and supercentenarians (> 110 years of age) (Murabito et al., 2012; Perls et al., 2007; Sebastiani and Perls, 2012). The current paradigm is lifespan is a complex polygenic trait, with the inheritance of multiple genes/variants with pleiotropic protective roles across several age-related diseases, and their interaction with environmental and lifestyle factors.

To identify genetic variation associated with human lifespan, research has focused on specific candidate genes and more recently common variant genome-wide association studies (GWAS) (Broer et al., 2015; Deelen et al., 2014; Deelen et al., 2019; Joshi et al., 2017; Pilling et al., 2017; Sebastiani et al., 2017; Zeng et al., 2016). While these past studies have identified multiple genetic variants associated with age, only one variant in apolipoprotein (*APOE*) is the most consistently replicated that passes genome-wide significance (p < 5×10^−08^) across multiple, independent cohorts and in meta-analyses (Partridge et al., 2018). The lack of replication in these studies may be due to small sample size, study-specific age cut-offs to define long-lived status, sex-specific genetic architecture of aging, and genetic and/or lifestyle heterogeneity among cohorts which may compromise meta-analyses. Moreover, the majority of published studies were performed on single candidate genes or common variant genome-wide genotype data, which limits the discovery of rare and novel age-related variants. In this study, we applied exome sequencing and exome-wide association analyses on ages of participants from large general populations and an extreme long-lived cohort to identify novel genetic variants associated with human aging.

## RESULTS

### Identifying genetic variants that change across strata of ages in the general population

The Regeneron Genetics Center (RGC) and Geisinger Health System (GHS) established the DiscovEHR study, consisting of GHS patients who participated in the MyCode Community Health Initiative. GHS serves a largely unselected population of >1.6 million participants from central Pennsylvania who are of predominantly European ancestry (Abul-Husn et al., 2016; Dewey et al., 2017; Dewey et al., 2016). As a discovery cohort, we analyzed exome sequence data from 58,470 participants of European descent (GHS60K) with a median age of 62 years (range 18-107 years) and median body mass index (BMI) of 30.3 kg/m^2^ (range 14.3-57.5 kg/m^2^) (**Figure 1A)**. GHS60K participants were female (n = 34,765; 59.4%) and male (n = 23,705; 40.6%), and majority were alive (n = 52,653; 90.1%) **(Figure 1B-D)**.

**Figure 1:**
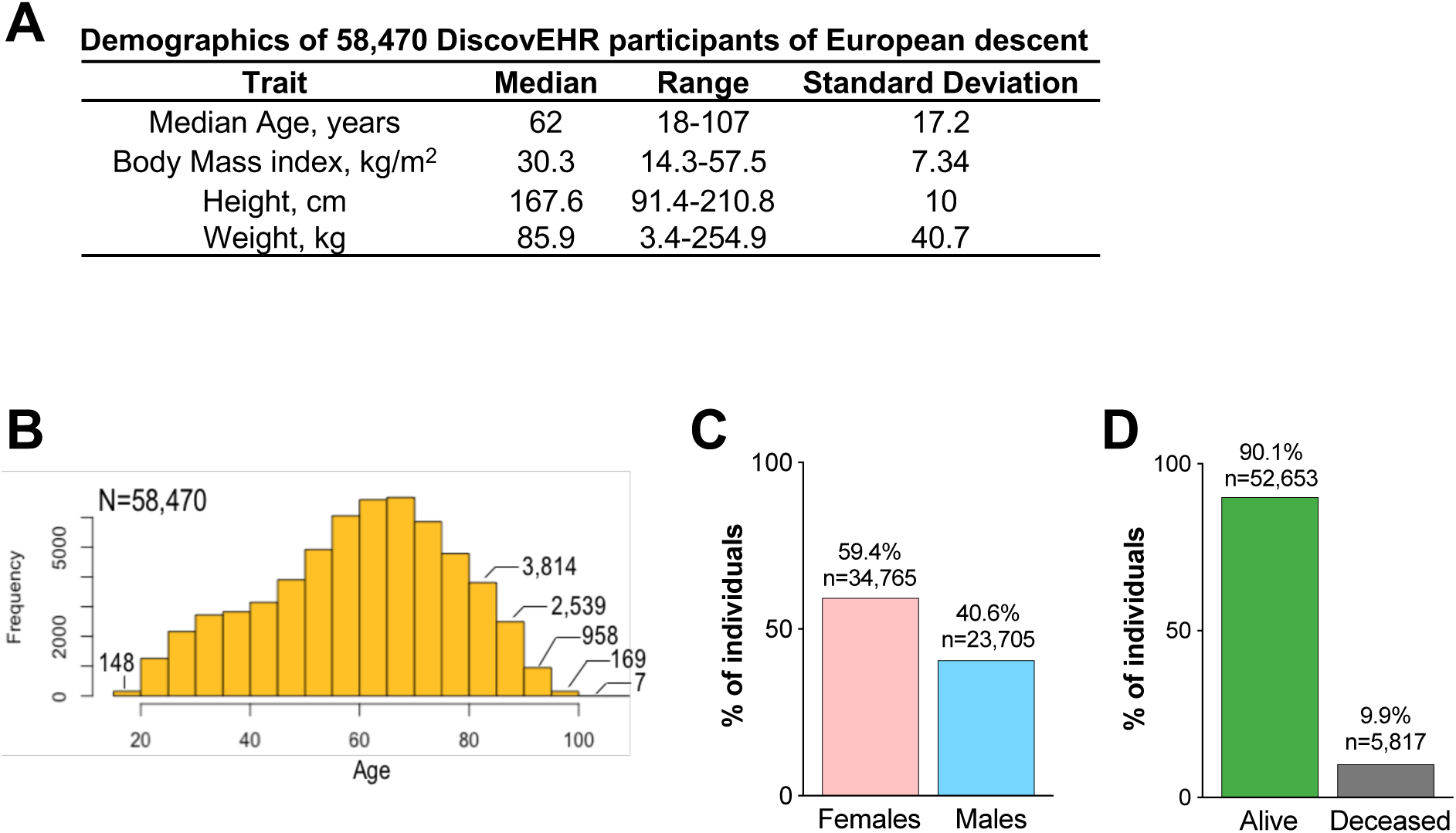
Demographics of DiscovEHR cohort. A) Table of median, ranges and standard deviation of age, body mass index, height and weight of 58,470 participants (GHS60K) of European descent from the DiscovEHR study. B) Age distribution of GHS60K; number of participants in bins 18-20 years and > 80years of age. C-D) Distribution of male/female and alive/deceased status for GHS60K participants.

To identify age-associated variants, we performed an ‘Age-ExWAS’ which entails a linear mixed model (LMM-BOLT) analysis using ‘Age at last encounter’ obtained from the electronic health record (EHR) as a quantitative trait, as shown in Figure 1B, under an additive model, which was adjusted for four principal components and sex as covariates **(Figure 2A)**. First, as proof-of-concept, we tested whether 73 previously published candidate age-related variants were also age-associated in our analysis (Partridge et al., 2018), of which 16 were present in GHS60K exome data **(Supplementary Table 1)**. We replicated *APOE* (rs429358), *APOE* (rs2075650) and *HFE* (rs1800562) variants, which were negatively associated with age and passed the Bonferroni adjusted p-value threshold (p = 3.10×10^−03^) **(Figure 2B)**. We next calculated the allelic frequencies of *APOE* haplotypes per age group, which revealed the common *APOEe4* haplotype, which is associated with Alzheimer’s disease and early mortality, decreases in frequency with age (p = 2.20×10^−12^). In contrast, the protective and rarer *APOEe2* haplotype increases in frequency with age (p = 9.90×10^−08^) **(Figure 2C)**.

**Figure 2:**
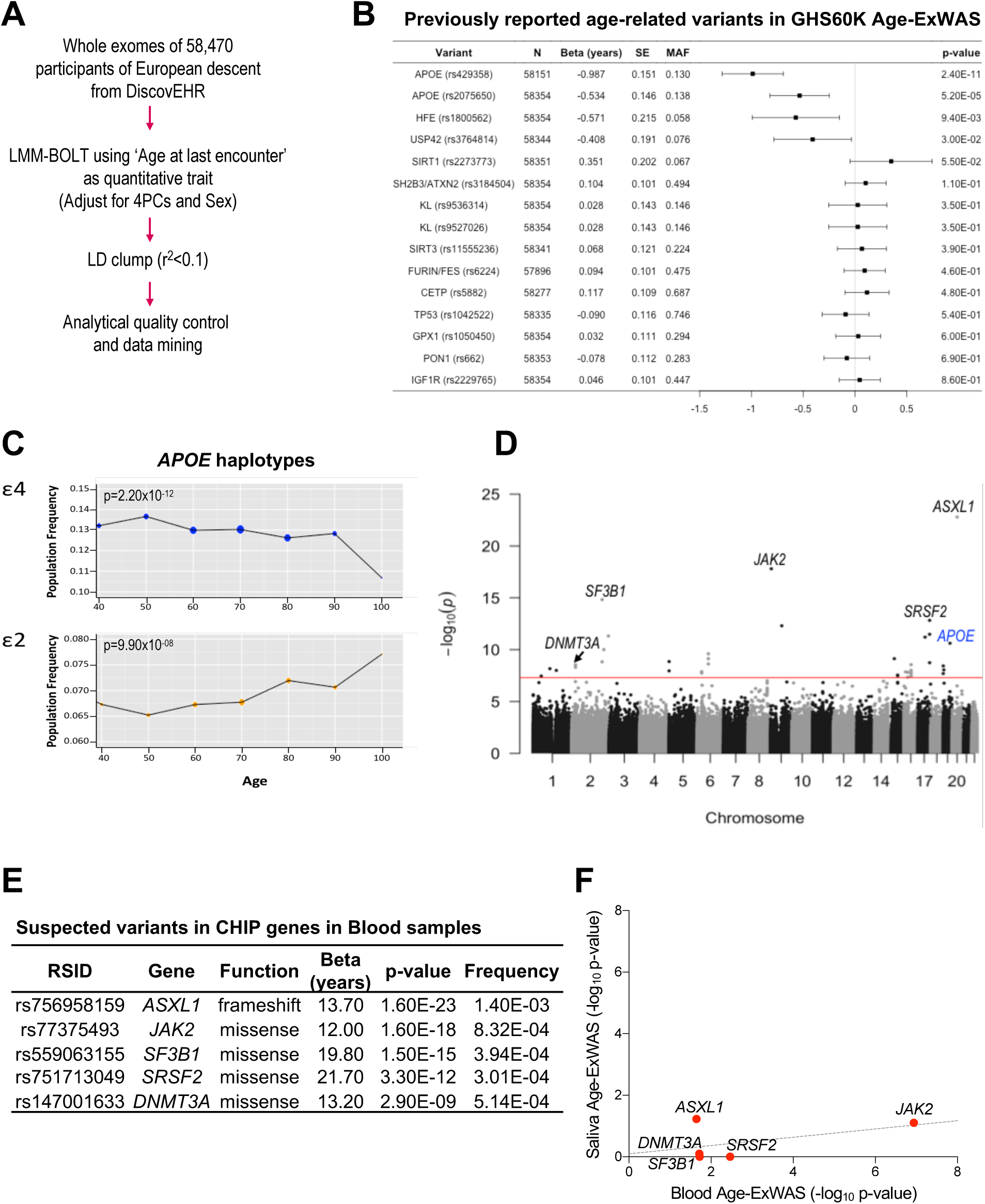
*APOE* and CHIP variants are top age-associated hits in Age-ExWAS. A) Linear mixed model (LMM-BOLT) analysis on exome sequence data using ‘Age at last encounter’ as the trait of interest under the additive model and normalized for sex and 4 principal components, followed by LD clumping (r^2^ < 0.1). Analytical quality control was performed using criteria defined in methods. B) Forest plot of reported age-related variants from published literature. Variant name, N, Beta (non-transformed), SE, MAF and p-value are shown. X-axis shows Beta in years. C) Allele frequencies of carriers of *APOEe*2 and *APOEe4* haplotypes plotted based on age deciles. D) Manhattan plot of Age-ExWAS (λ = 1.1) in GHS60K. Associations - log10(p-value) for each genome-wide SNP (y-axis) by chromosomal position (x-axis). Red line indicates the threshold for genome-wide statistical significance (p = 5×10^−08^). Variants reported as clonal hematopoiesis of indeterminate potential (CHIP) and *APOE* are shown in black and blue, respectively. E) Table of top 5 suspected CHIP variants where RSID, gene name, prediction function, Beta (non-transformed), p-value and MAF. F) Age-ExWAS on exome sequence data of DNA extracted from 8,102 blood or 8,102 saliva samples. The top suspected CHIP variants are shown in the scatterplot where -log10(p-value) of Blood Age-ExWAS (x-axis) and Saliva (Age-ExWAS) y-axis. SE = standard error; MAF = minor allele frequency.

Age-ExWAS revealed variants in genes known to be associated with clonal hematopoiesis of indeterminate potential (CHIP), which exhibited the most significant p-values and were positively associated with age in our analysis. These included variants with predicted deleterious functions in *ASXL1* (rs756958159), *JAK2* (rs77375493), *SF3B1* (rs559063155), *SRSF2* (rs751713049) and *DNMT3A* (rs147001633) **(Figure 2D-E)**. CHIP variants are defined as the accumulation of age-related somatic mutations resulting in clonal expansion of hematopoietic cells in the absence of hematological malignancies clinical phenotypes (Steensma et al., 2015). To confirm whether these suspected CHIP variants are indeed blood-borne somatic mutations that were designated as “heterozygotes” with our genotype calling algorithm, we performed a comparative Age-ExWAS on exome sequence data derived from DNA extracted from saliva (n = 8,102; median age = 51.6 years) or a matched sample size from blood (n = 8,102; median age = 55.2 years) of the DiscovEHR cohort, which was adjusted for four principal components and sex. Our analysis revealed the five suspected CHIP variants from GHS60K Age-ExWAS analysis were more significantly associated with age from blood samples, in contrast to saliva **(Figure 2F)**. Collectively, this data suggest that these top age-associated variants are likely somatic CHIP variants from blood.

In addition to *APOE* and suspected CHIP variants, Age-ExWAS in the GHS60K discovery cohort identified 18 additional variants that exceeded genome-wide significance (p < 5×10^−08^) **(Figure 3A)**. To validate our findings, we performed Age-ExWAS analysis on two additional replication cohorts, including GHS30K (n = 28,930; median age = 54 years) and UPENN biobank (n = 8,209; median age = 68 years) **(Figure 3B; Supplementary Table 2)**. Interestingly, we observed a rare variant in *DISP2* (rs183775254) which was validated with nominal significance and directional consistency in both replication cohorts (Meta-analysis: p = 2.80×10^−10^; Beta = −12 years; MAF = 3.50×10^−04^) **(Figure 3C)**. While we observed additional variants in/near *DISP2*, the intronic *DISP2* (rs183775254) SNP was the top age-associated variant in our analyses. Gene burden tests did not show any additional associations of *DISP2* with age **(Supplementary Table 3)**. This data indicates the specific *DISP2* rs183775254 variant, which is located within the first intron of *DISP2* and decreases in allele frequency with increasing age, as the top age-related and replicable signal in our analysis.

**Figure 3:**
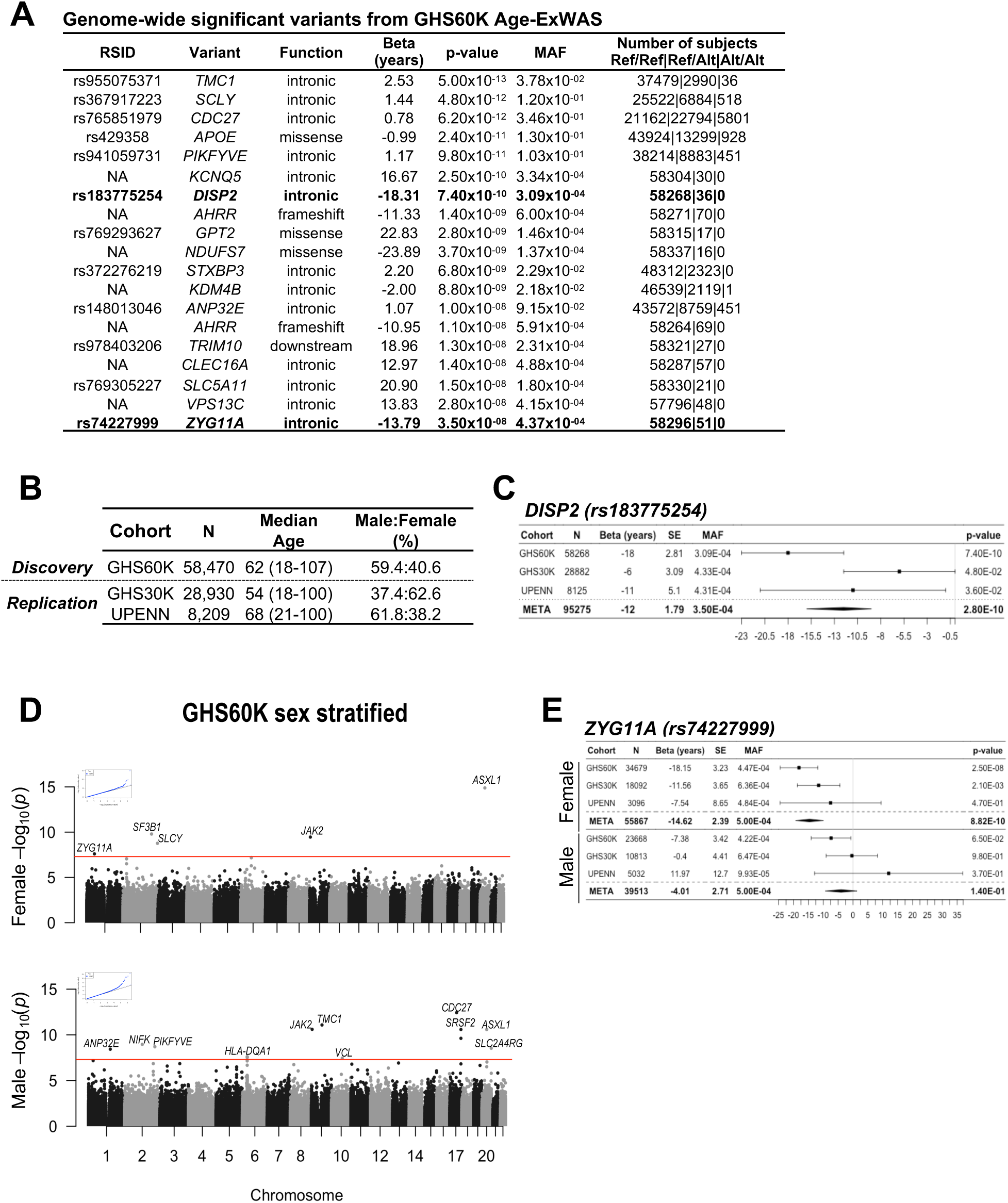
*DISP2* and *ZYG11A* as age-associated variants in general populations. A) Table of age-related variants from GHS60K Age-ExWAS analysis that passed genome-wide significance (p < 5×10^−08^). RSID, gene name, predicted function, untransformed Beta, p-value, MAF and number of carriers are shown. B) Demographics of replication cohorts of European descent including 28,930 additional participants from GHS30K and 8,209 participants from UPENN Biobank. Total N, median age and male-to-female ratio are shown. C) Forrest plot analysis showing Age-ExWAS results of *DISP2* variant in GHS60K, GHS30K and UPENN and meta-analysis. Non-transformed Beta and standard error (SE) in years, direction and p-values for each cohort are shown. D) Manhattan plot of female-only (λ = 1.07) and male-only (λ = 1.05) Age-ExWAS in GHS60K. Associations -log10(p-value) for each genome-wide SNP (y-axis) by chromosomal position (x-axis) are shown. Red line indicates the threshold for genome-wide statistical significance (p = 5×10^−08^). Top variants that passed genome-wide-significance are labeled. E) Forrest plots of *ZYG11A* in female and male GHS60K, GHS30K, UPENN and meta-analysis. SE = standard error; MAF = minor allele frequency.

As this *DISP2* variant is negatively associated with age (−12 years in meta-analysis), we next characterized demographics of carriers of this SNP. As expected, the 36 carriers of this variant in GHS60K are younger (median age = 38.8 years; range = 20-87), as compared to 56,174 non-carriers (median age = 61.4 years; range = 18-105). We performed phenome-wide association studies (Phe-WAS) using GHS and UK Biobank (UKB) phenotype data of participants of European ancestry, which showed no International Statistical Classification of Diseases and Related Health Problems (ICD-10) codes significantly associated with *DISP2* rs183775254 **(Supplementary Table 4)**. To explore the clinical significance this SNP, we analyzed electronic health records of 36 carriers of rs183775254 and age/sex-matched non-carriers to identify ICD-10 codes diagnosed these patients. Interestingly, total burden of ICD-10 codes was significantly higher (p = 4.0×10^−03^) in carriers (2,104 total; median = 51 ICD-10 codes/patient) versus non-carriers (1,253 total; median = 30 ICD-10 codes/patient), which comprised of 16 disease categories **(Supplementary Table 5)**.

*DISP2* is the human homologue of the Dispatched gene in Drosophila, which acts as a transporter-like membrane protein that releases cholesterol-modified sonic hedgehog (SHH) proteins to trigger long-range SHH signaling with roles in early development and embryonic patterning. As the specific *DISP2* rs183775254 SNP is intronic, we next sought to map whether this was located in a transcriptional regulatory region by aligning the position of this variant to publicly available regulatory datasets, including DNAse I hypersensitivity, which marks open transcription factor-accessible chromatin, and H3K27Ac chromatin immuno-precipitation sequencing (ChIP-seq) data, which identifies enhancer elements. Notably, we observed this variant is located within an open chromatin region enriched in H3K27Ac marks specific to early human fetal neural stem cells and absent in other cell types, consistent with high *DISP2* levels in brain tissue **(Supplementary Figure 1A-B)**. Mapping of transcription factor binding sites showed this region is enriched in multiple enhancer-related transcription factors, including MAX, POLR2A, EP300, JUND, RAD21 and CTCF, indicating this variant may be located in a transcriptionally active regulatory hotspot. Further analysis of expression quantitative trait loci (eQTL) data from GTEx in brain tissue did not show associations with *DISP2* rs183775254. However, four other *DISP2* variants were associated with cis eQTL in brain tissue with positive effects on *DISP2* expression, which include rs71472433 (brain caudate basal ganglia; p = 8.99×10^−28^; Beta = 0.65), rs56221586 (putamen basal ganglia; p = 2.17×10^−24^; Beta = 0.71), rs12913300 (brain nucleus accumbens basal ganglia; p = 7.30×10^−20^; Beta = 0.52), and rs56221586 (cortex; p = 5.88×10^−11^; Beta = 0.32). Collectively, these observations suggest a potential role of *DISP2* and SHH signaling in early brain development and aging.

As multiple studies suggest a gender specific component of aging, we next performed a sex-stratified analysis, in which Age-ExWAS analysis was applied to GHS60K females (n = 34,756; median age = 58 years) and males (n = 23,705; median age = 65 years). These analyses revealed 5 and 11 variants that passed genome-wide significance in GHS60K females and males, respectively **(Figure 3D; Supplementary Table 2)**. Variant lookup in sex-stratified GHS30K and UPENN Age-ExWAS analyses showed that CHIP variants, including *JAK2, SRSF2, ASXL1*, replicated as age-associated variants in both males and females. In addition, we identified an intronic variant in *ZYG11A* (rs74227999) significantly associated and negatively correlated with age in females, but not in males **(Figure 3E)**. Further meta analyses strengthened this sex-specific associations in females (p = 8.82×10^−10^; Beta = −14.62 years; MAF = 5.00×10^−04^) as compared to males (p = 1.40×10^−01^; Beta = −4.01; MAF = 5.00×10^−04^). Gene burden test analyses for *ZYG11A* did not show any significant associations, pointing to *ZYG11A* (rs74227999) as the top replicable and age-related variant in our sex-stratified analysis **(Supplementary Table 3)**.

We next characterized 51 carriers of the *ZYG11A* rs74227999 variant in GHS60K, which comprised of 32 female and 19 males. We observed female carriers (median age = 37 years; range = 20.3-73.1) were younger than male carriers (median age = 59.7 years; range = 20.9-89.7). Phe-WAS analysis showed no obvious ICD-10 code association with this variant **(Supplementary Table 4)**. Notably, we observed a significantly higher (p = 2.39×10^−06^) burden of diagnosed ICD-10 codes in the 51 carriers (4,082 total; median = 54 ICD-10 codes/patient), as compared to 51 age/sex-matched non-carriers (1,983 total; median = 37 ICD-10 codes/patient), which are classified by disease categories **(Supplementary Table 5)**.

*ZYG11A* is a member of cell cycle regulators which has roles in driving cellular proliferation. Epigenetic mapping showed the intronic *ZYG11A* rs74227999 variant is located within a euchromatic region with an active enhancer specific to GM12878, a lymphoblastoid cell line derived from blood of a female donor of European decent and enriched for multiple enhancer-related transcription factor binding sites. While *ZYG11A* is highly expressed in testis tissues **(Supplementary Figure 2A-B)**, we did not observe significant association of rs74227999 with eQTL in germinal tissues. However, we observed one variant in *ZYG11A* rs534070 (p = 4.26×10^− 13^; Beta = 0.29) that was positively associated with cis eQTL is testis. Taken together, these observations indicate rare variants in non-coding regions within *DISP2* and *ZYG11A* may be located within transcriptional regulatory regions, which may affect expression of these or nearby genes.

### Exome- and genome-wide association analysis of long-lived case (LLI)-Control cohorts

There is a general discrepancy in defining the age threshold of long-lived status. Studies calculating the relative ‘risk’ of siblings living to the ages of their oldest-living sibling showed that the risks increase significantly at extreme ages (> 100 years) (Gudmundsson et al., 2000; Kerber et al., 2001; Sebastiani et al., 2016). These studies also indicate that the difference in ‘relative risk’ of living to 90 or 95 years is very marginal, which suggests limited heritability of longevity at these ages. Moreover, multiple extreme long-lived individuals are classified as ≥ 95 years of age, regardless of gender (Ayers et al., 2017; Deluty et al., 2015; Gubbi et al., 2017; Perice et al., 2016). As females generally live to older ages, we calculated and applied a sex-specific age threshold for long-lived status. Based on the 2015 Mortality Data from the National Vital Statistics, Center for Disease Control of Americans of European ancestry, which records the total number and ages of death per 100,000 individuals (Murphy et al., 2017), we defined 95 years as a cutoff for long-lived status in females, which represented 11.8% of female deaths in 2015 **(Figure 4A)**. In males, 11.8% of deaths in 2015 occurred at ≥ 93 years of age. We extended this analysis to include mortality data from 1997-2015, in which we calculated that 95 years for female is equivalent to a median age of 92 years for males **(Figure 4B)**. Thus, in our subsequent analyses, we classified as a long-lived-individual (LLI), ≥ 95 years for females and ≥ 92 years for males.

**Figure 4:**
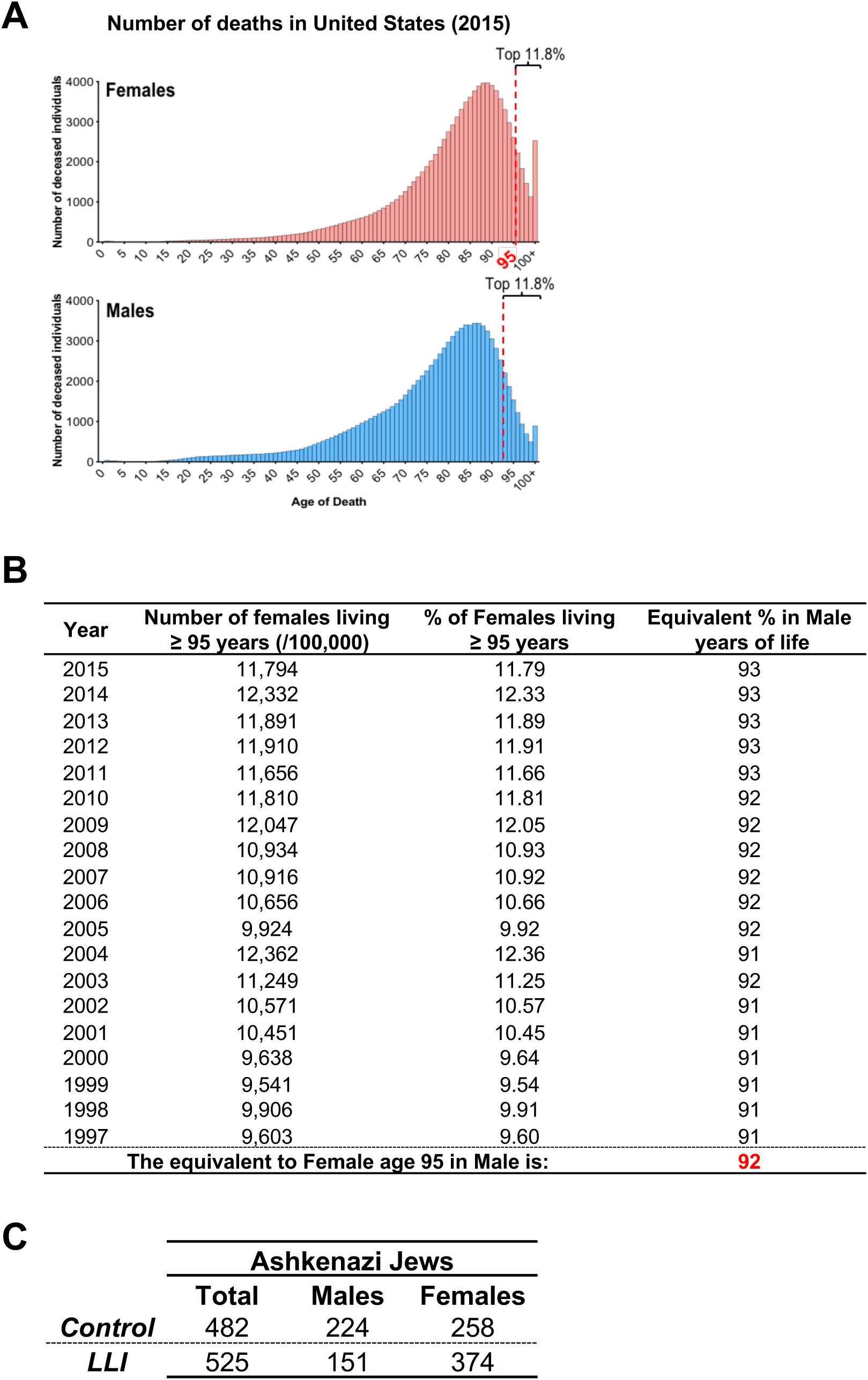
Determination of sex-specific age threshold of long-lived status. A) Distribution of total number and ages of deaths per 100,000 individuals of European descent in the United States in 2015. B) Mortality data (1997-2015) showing 95 years in female is equivalent to 92 years in males. C) Table showing total N, median ages and sex distribution of controls and long-lived individuals (LLI) of Ashkenazi Jewish descent.

We exome sequenced a large collection of LLI of Ashkenazi Jews, a founder population with high degree of endogamy that exhibits longevity phenotypes and decreased incidence of age-related pathologies (Atzmon et al., 2004). Based on our criteria, there were 525 LLI of Ashkenazi Jewish descent (151 males, 374 females) and 482 controls (224 males, 258 females) **(Figure 4C)**, where both the father and mother of the controls passed away before 92 or 95 years of age, respectively. To identify rare and common variants associated with extreme ages, we merged exome sequence data and genome-wide chip genotype data imputed to the 1000 Genomes reference panel and performed Case-Control analyses (SAIGE – refer to methods) using Control and LLI groups as a binary trait, followed by further adjusting for sex as a covariate **(Figure 5A)**. As positive control, we calculated frequencies of *APOE* haplotypes in Ashkenazi Jews, which revealed LLIs were enriched for the protective *APOEe2* allele and depleted for the *APOEe4* risk allele, as compared to controls **(Figure 5B)**. Moreover, we assessed for association with longevity 28 previously reported candidate variants (Partridge et al., 2018; Singh et al., 2019) and identified variants in *APOE* (rs6857, rs4420638) and *TMTC2* (rs7976168) that passed a Bonferroni-adjusted statistical significance threshold of (p = 1.70×10^−03^) **(Figure 5C; Supplementary Table 1)**.

**Figure 5:**
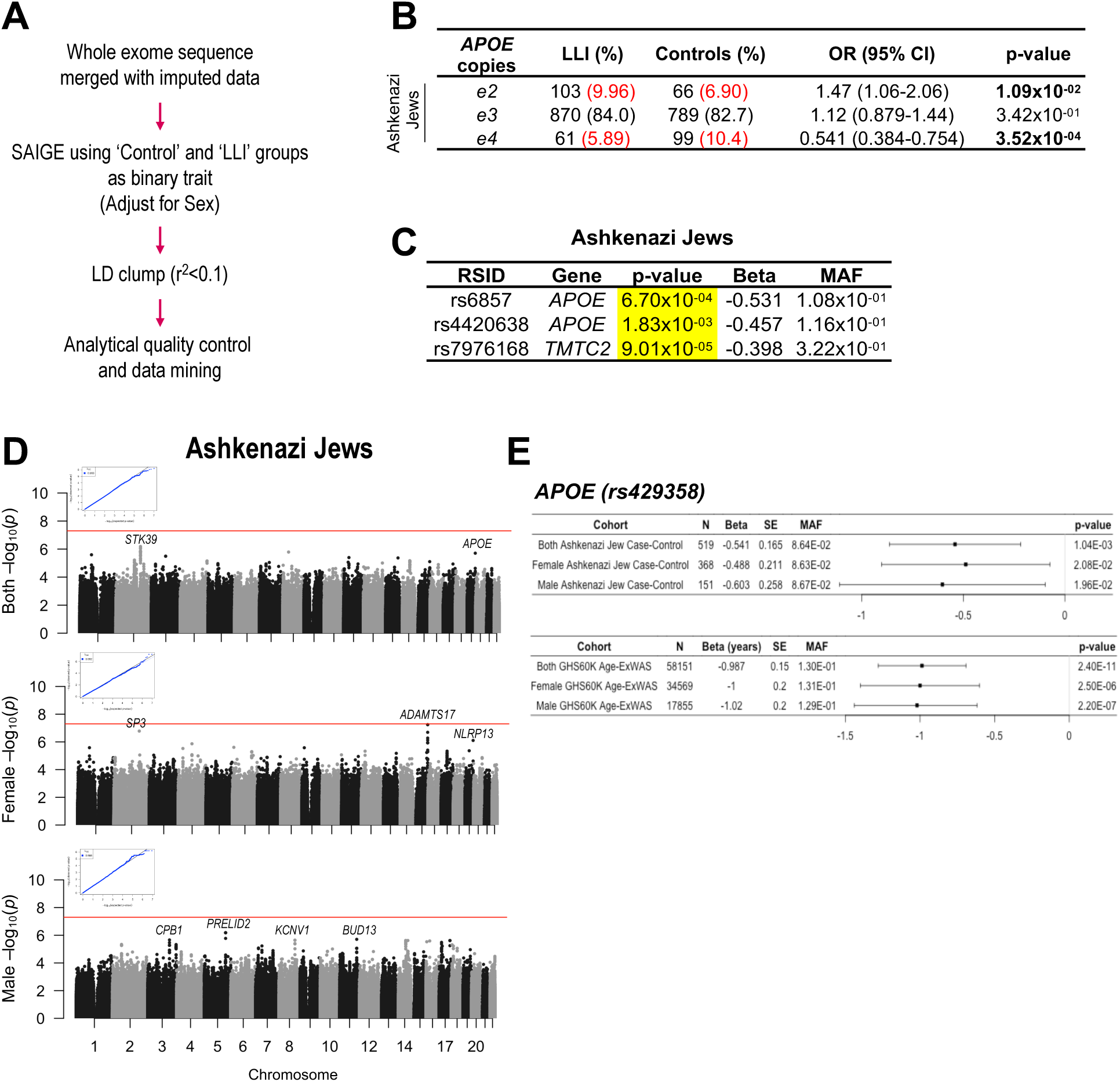
Case-control analyses in Ashkenazi Jews long-lived cohort. A) Case-Control analysis using SAIGE was performed on exome sequence data merged with genotype data imputed to 1000 Genome reference panel using Control and LLI groups as a binary trait, followed by further adjusting for sex as covariate, followed by LD clumping (r^2^ < 0.1). Analytical quality control was performed using criteria defined in methods. B) Table showing frequencies, odd ratio (OR) and p-values of *APOEe2-4* haplotypes in LLIs and controls. Percentage of individuals carrying the given *APOE* haplotype copies shown in red. C) Of 28 published positive control SNPs (Supplementary Table 1), variants that passed Bonferroni adjusted p-values (p = 1.70×10^−03^) are highlighted in yellow. D) Manhattan plots for Ashkenazi Jew cohort both sexes combined, female-only and male-only showing associations of -log10(p-value) for each genome-wide SNP (y-axis) by chromosomal position (x-axis). Red line indicates the threshold for genome-wide statistical significance (p = 5×10^−08^). Top-age-associated variants are labelled with nearest gene. QQ plots depicting minor genomic inflations are shown. E) Forest plot shows variant in *APOE* rs429358 in both sexes, female and males only in Ashkenazi Jews (top) and GHS60K Age-ExWAS (bottom). OR = odds ratio; SE = standard error; MAF = minor allele frequency.

In exome/genome-wide Case-Control analysis in Ashkenazi Jews, the top associated variant for both sexes combined was in *STK39* (rs7594207; intronic; p = 6.55×10^−07^; Beta = 0.521). In females only, the top variants were *ADAMTS17* (rs73484155; intronic; p = 5.70×10^−08^; Beta = −0.981) and *SP3* (rs12613192; intergenic; p = 1.66×10^−07^; Beta = −1.70) **(Figure 5D; Supplementary Table 6)**. In contrast, the top (p < 1×10^−05^) associated variants in male Case-Control analyses was in *PRELID2* (rs1982070; intergenic; p = 6.55×10^−07^; Beta = 0.28). Interestingly, of the top signals, only the variant in *APOEe4* rs429358 passed the Bonferroni adjusted p-value threshold with directional consistency in both Ashkenazi Jew case-control analyses (p = 1.04×10^−03^), as well as GHS60K Age-ExWAS analyses (p = 2.40×10^−11^) **(Figure 5E)**. Taken together, our results suggest both cohort and sex-specific variants associated with extreme ages worthy of follow up in an expanded sample of Ashkenazi Jewish, as well as other extreme long-lived cohorts.

## DISCUSSION

The discovery of new genetic variants associated with human aging is rapidly increasing due to advancement and lowered costs of genome-wide genotyping and next-generation sequencing technologies. In this study, we performed Age-ExWAS analysis on exome sequence data from large general population cohorts. We identified variants in genes involved in age-related clonal hematopoiesis as top age-associated candidates. Somatic variants in CHIP genes are detected in ∼1% of individuals < 50 years of age, but increase rapidly in frequency with age, where CHIP mutations are detected in ∼10% and ∼20% in persons aged 70 and 90 years of age, respectively (Dorsheimer et al., 2019; Genovese et al., 2014; Jaiswal et al., 2014; Zink et al., 2017). High frequencies of CHIP variants have been reported to be associated with hematologic malignancies as well as atherosclerosis, although whether the latter is independent of age requires further investigation. The most commonly recurrently mutated CHIP genes are epigenetic and chromatin modifying enzymes *DNMT3A, ASXL1* and *TET2*. Indeed, bone-marrow transplant of *TET2*-deficient hematopoietic stem cells was sufficient to accelerate clonal expansion and increase atherosclerotic plaque formation in immunodeficient mouse models, suggesting a causal role of CHIP variants (Fuster et al., 2017). Interestingly, our analyses revealed CHIP variants also present in long-lived persons of Ashkenazi Jewish descent, without history of cancer or cardiovascular diseases (data not shown). Whether total numbers of CHIP variants or a specific combination may act as a biomarker of healthy versus degenerative aging remains to be studied. Additionally, longitudinal studies focused on transcriptional and epigenetic profiling of blood cells with an increasing number of CHIP variants may shed light into the specific cellular states required for CHIP-mediated pathogenesis.

Our analysis identified a rare age-associated variant within *DISP2*, in which epigenetic mapping revealed to be located within an enhancer specific to primitive neural cells. *DISP2* is an upstream effector of SHH signaling with roles in embryonic development and is highly expressed neural tissue (Hall et al., 2019). In the absence of Disp, hedgehog ligands fail to release from donor cells, resulting in impaired early neural and embryonic development (Burke et al., 1999; Ma et al., 2002). In zebrafish, *disp1* transcripts are detected along the skeletal rod and musculature, whereas *disp2* is present in the telencephalon and hindbrain (Nakano et al., 2004). *In vivo* antisense oligonucleotide-mediated silencing of *disp1*, but not *disp2*, resulted in disrupted downstream SHH signaling, suggesting alternate downstream mechanisms may be regulated by *disp2* (Li et al., 2008). Our studies suggest a potential role of *DISP2* in aging, perhaps with relevance to the brain and nervous system, which will require further validation in other populations and functional assessment in animal and cell systems.

We also identified a rare *ZYG11A* variant to be significantly associated with age in females that is located within an enhancer element in lymphoblastic cells. In C.elegans, *zyg11* is implicated in meiotic progression and early embryogenesis and exhibits a maternal effect lethal phenotype, in which offspring of homozygous mutant mothers are embryonic lethal (Carter et al., 1990). Intriguingly, *ZYG11A* is highly expressed in testis, which indicates potential sex-specific role of the *ZYG11A* rs74227999 variant in females, whereas other variants in this gene may function in male reproductive development. Additional functional validation in sex-stratified models will be needed to test the gender specific roles of *ZYG11A* variants. High levels of *ZYG11A* are associated with advanced lung adenocarcinoma and drives cellular proliferation in part via upregulation of cyclin E to promote tumorigenesis (Husni et al., 2019; Wang et al., 2016). Recent studies implicated *ZYG11A* as downstream of the IGF-1 pathway in p53 wild-type cancer cells (Achlaug et al., 2019). As it is increasingly evident that suppression of IGF signaling is connected to healthy aging and longevity (Harrison et al., 2009; Selman et al., 2009), our data suggests that *ZYG11A* may play a role in aging, potentially through modulation of IGF signaling.

Consistent with negative associations of these variants with age, analyses of clinical data revealed significantly higher burdens of ICD-10 codes in *DISP2* rs183775254 and *ZYG11A* rs74227999 carriers, as compared to age and sex matched non-carrier controls. Specifically, our data suggests that carriers of these variants exhibit higher diagnoses of ICD-10 codes related to infections, neoplasms, neurodevelopmental disorders and diseases of the vasculature, nervous system, circulatory system, respiratory system, skin and subcutaneous tissue, musculoskeletal system and connective tissues and genitourinary system. The precise biological and genetic mechanisms by which these variants increase disease susceptibility and diagnoses are actively being studied.

Similar with prior longevity studies (Kulminski et al., 2016; Schachter et al., 1994; Sebastiani et al., 2019), our Case-Control association analyses showed association with increased and decreased frequencies of *APOEe2* and *APOEe4* haplotypes, respectively, in LLI as compared to controls. As with many prior GWAS studies (Partridge et al., 2018), our analysis revealed *APOE* as the only locus that was age-associated and replicated in our studies. Our Case-Control analyses also identified cohort-specific variants suggestively associated with extreme ages. It is important to highlight that while of European ancestry, the Ashkenazi Jews are a founder population with distinct allelic architecture. As the prevalence to living to such extreme long-lived ages is estimated to be ∼1/10,000 (Perls et al., 1999), additional efforts will be required to recruit larger cohorts of long-lived individuals for exome profiling to replicate our findings and to identify additional rare longevity-related variants.

In addition to general population regression analyses and case-control analyses in extreme-aged cohorts, an alternative approach to identify aging-related variants is using parental ages of death regressed on offspring genotypes. Recent parental lifespan studies from large populations including the UK Biobank and AncestryDNA genotype data have identified *APOE, LPA, CHRNA3/5, LDLR, SH2B3/ATXN2, CDKN2B-AS1* and other study-specific variants as associated with parental ages (Joshi et al., 2017; Pilling et al., 2017; Timmers et al., 2019; Wright et al., 2019). In parallel with our case-control analyses on extreme-aged cohorts, it will be of high interest to extend these analyses to include parental lifespan analyses on exome sequence data to identify rare, coding variants associated with aging.

In summary, our study applied exome sequencing in large patient populations and LLI Case-Control cohorts and identified several candidate gene variants associated with age and/or longevity. Further studies will be necessary for further replication and functional validation of genes/variants associated with aging to gain mechanistic insights into aging biology and to inform interventions to promote healthy aging.

## AUTHOR CONTRIBUTIONS

P.S-C and A.R.S conceived the project and wrote the manuscript. All analyses were performed by P.S-C with assistance from N.G, C.G, C.V.V.H, B.Y, A.M, A.H.L, C.O and DL. Exome sequencing and variant calling were performed under the supervision of J.D.O, J.D.R, and A.B. A.E and D.G provided scientific input. DNA for exome sequencing and electronic medical records were generously provided by D.J.C, D.H.L, D.R, M.D.R, S.M.D, S.M and N.B.

## EXPERIMENTAL PROCEDURES

### Study design and participants

Age-ExWAS analysis was performed using genomic DNA samples and data from four cohorts, including two DiscovEHR study populations from the MyCode Community Health Initiative of Geisinger Health System (GHS). The GHS discovery cohort (GHS60K) consisted of 58,470 persons of European ancestry who were recruited from outpatient primary care and specialty clinics. Replication cohorts consisted of 28,930 additional persons of European ancestry from the DiscovEHR study (GHS30K) and 8,209 persons of European ancestry from the University of Pennsylvania (UPENN) Biobank. Our case-control analyses consisted of 1,007 persons of Ashkenazi Jewish descent, recruited by Albert Einstein College of Medicine (Barzilai et al., 2003).

### DNA Sample Preparation and sequencing

DNA sample preparation and exome sequencing for participants were performed at the Regeneron Genetics Center, as previously described (Abul-Husn et al., 2016; Dewey et al., 2017; Dewey et al., 2016). Briefly, exome capture was performed using a custom reagent kit from Kapa Biosystems using a fully-automated approach developed at the Regeneron Genetics Center. A unique 6 base pair barcode was added to each DNA fragment during library preparation to facilitate multiplexed exome capture and sequencing. Equal amounts of sample were pooled prior to exome capture with NimbleGen probes on SeqCap VCRome or IDT xGen platforms. Multiplexed samples were sequenced using 75 bp paired-end sequencing on an Illumina v4 HiSeq 2500 to a coverage depth sufficient to provide greater than 20x haploid read depth. Raw sequence data from each Illumina Hiseq 2500 run were uploaded to DNAnexus (Reid et al., 2014) for sequence read alignment and variant identification. Raw sequence data were converted from BCL files to sample-specific FASTQ-files, which were aligned to the human reference build GRCh37.p13 with BWA-mem (Li and Durbin, 2009). Single nucleotide variants (SNV) and insertion/deletion (INDEL) sequence variants were identified using the Genome Analysis Toolkit (McKenna et al., 2010).

### Exome-wide association study and analytical quality control

We utilized BOLT-LMM v2.2 to test for associations between variants and age for DiscovEHR and UPENN exome sequence data, which uses the mixed models of association approach (Loh et al., 2015). Genetic relatedness matrix (GRM), which captures population structure from ancestry and relatedness, was included as a random-effects covariate. The GRM was constructed from 39,858 non-MHC markers, with no greater than 1% genotype missingness, and with minor allele frequency > 0.1%. Logistic regression analyses adjusted for sex and the first four principal components of ancestry. Individual study results were meta-analyzed using METAL (Willer et al., 2010) on plink 1.9. Case-control analyses on Ashkenazi Jewish cohorts were performed using Scalable and Accurate Implementation of GEneralized mixed model (SAIGE) to control for sample relatedness (Zhou et al., 2018), and adjusted for sex as covariate (except in analyses stratified by sex). Variants were filtered using a Minor Allele Count of ≥ 5. Following this, SNPs were filtered using Quality Depth of ≥ 3, Read Depth of ≥ 7 and Allele Balance of ≥ 15/85, whereas INDEL were filtered using Quality Depth of ≥ 5, Read Depth of ≥ 10 and Allele Balance of ≥ 20/80.

### Imputation

Ashkenazi Jewish genome-wide genotype data (Illumina GSA array) of European participants were converted to hg19 format, duplicate SNPs removed, and quality controlled by excluding variants with high missingness (> 10%), high deviations from Hardy-Weinberg equilibrium (1.00×10^−15^) and minor allele frequency < 1%. Variants were imputed to HRC-1000G reference panel using Michigan Imputation Server. Imputed variants with info score > 0.3 were kept, lifted-over to hg38 format and merged with exome sequence data. For overlapping variants, those from exome data were selected.

### Epigenetic mapping of regulatory regions

DNase I hypersensitive site, histone marks and transcription factor binding sites were downloaded from ENCODE cell line public data. H3K27Ac ChIP-seq data on human neural stem cell was downloaded from previously published studies (Sin-Chan et al., 2019).

### Calculation of *APOE* haplotype copies

To calculate number of copies of *APOE* haplotypes in long-lived cohorts, the rs429358 (Cys130Arg) and rs7412 (Arg176Cys) variants were extracted from exome sequence data. Based on the specific combinations, we determined the *APOE* haplotypes: *e2* (Cys130/Cys176), *e3* (Cys130/Arg176) and *e4* (Arg130/Arg176). We restricted our analysis on individuals with *e2/e2, e2/e3, e3/e3, e3/e4* and *e4/e4* genotypes, resulting in 475 controls and 517 LLI of Ashkenazi Jewish descent. The p-value, odds ratio and confidence intervals were calculated using logistic regression, which was normalized for Sex as covariate.

## SUPPLEMENTARY FIGURE LEGENDS

**Supplementary Figure 1:**
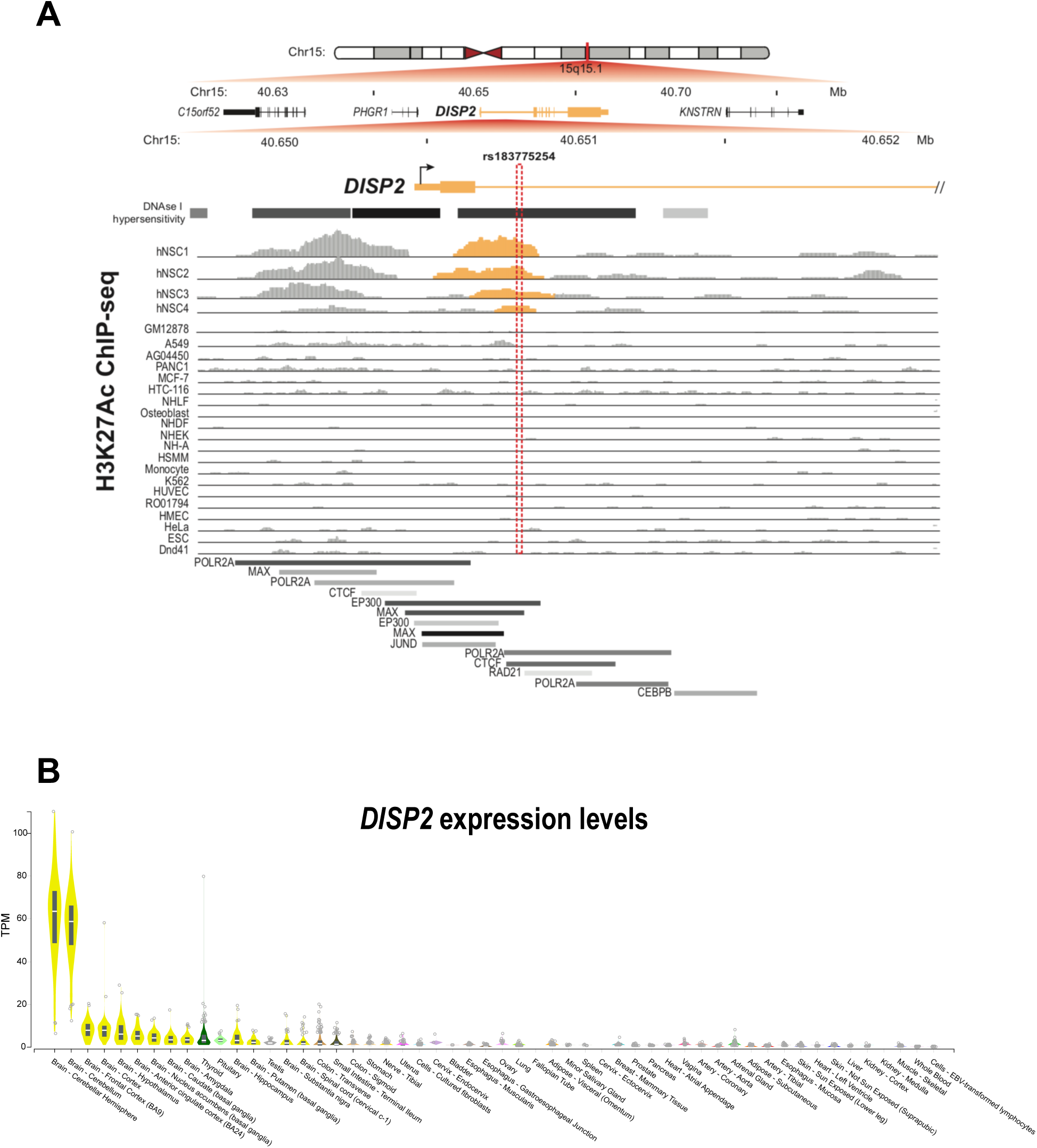
Enhancer mapping and tissue expression of *DISP2*. A) Schematic map of *DISP2* (chr15:40,649,698-40,652,202) relative to UCSC hg19 RefSeq annotation and ENCODE tracks. Variant of interest is highlighted in the red box and shown relative to ENCODE DNAse I hypersensitive sites, H3K27Ac-ChIP-seq from fetal human neural stem cells and ENCODE cell lines and ENCODE ChIP-seq map of enhancer-related transcription factor binding sites. B) Expression levels of *DISP2* from Genotype-Tissue Expression (GTEx) database.

**Supplementary Figure 2:**
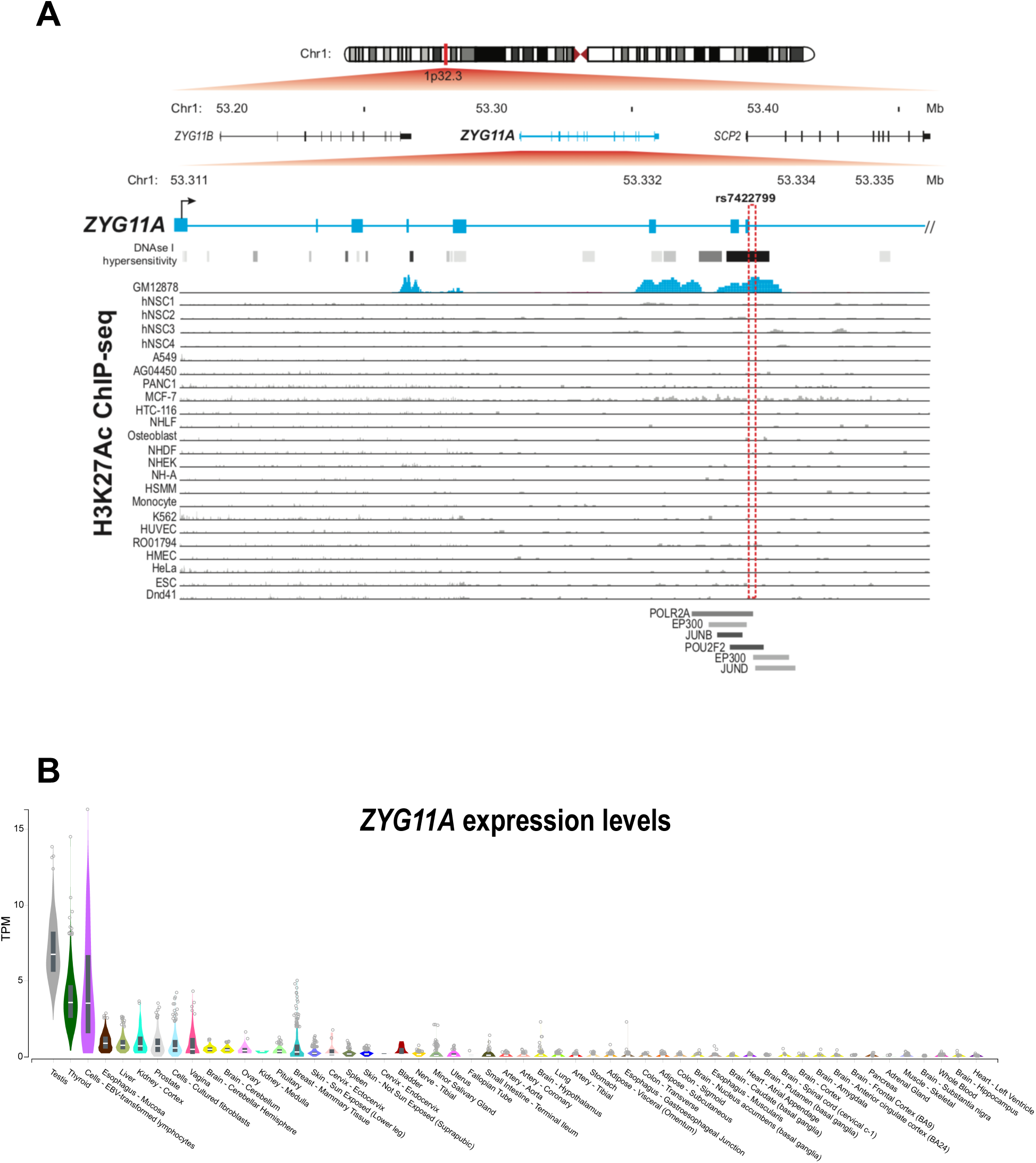
Enhancer mapping and tissue expression of *ZYG11A*. A) Schematic map of *ZYG11A* (chr1:53,310,672-53,335,672) relative to UCSC hg19 RefSeq annotation and ENCODE tracks. Variant of interest is highlighted in the red box and shown relative to ENCODE DNAse I hypersensitive sites, H3K27Ac-ChIP-seq from fetal human neural stem cells and ENCODE cell lines and ENCODE ChIP-seq map of enhancer-related transcription factor binding sites. B) Expression levels of *ZYG11A* from Genotype-Tissue Expression (GTEx) database.

## SUPPLEMENTARY TABLES

**Table 1** – Positive control variants and lookup in analyses

**Table 2** – Genome-wide significant variants from GHS60K Age-ExWAS analysis and lookup replication cohorts

**Table 3** – *DISP2* and *ZYG11A* variants and Burden test results in GHS60K Age-ExWAS

**Table 4** - Phenome-wide association study (Phe-WAS) analysis of age-related variants

**Table 5** – Distribution of disease traits in *DISP2* and *ZYG11A* carriers

**Table 6** – Ashkenazi Jew Case-Control top hits (p < 1×10^−05^) and lookup in GHS60K Age-ExWAS analyses

